# Antimicrobial resistant *Klebsiella pneumoniae* carriage and infection in specialized geriatric care wards linked to acquisition in the referring hospital

**DOI:** 10.1101/218461

**Authors:** Claire L. Gorrie, Mirjana Mirceta, Ryan R. Wick, Louise M. Judd, Kelly L. Wyres, Nicholas R. Thomson, Richard A. Strugnell, Nigel F. Pratt, Jill S. Garlick, Kerrie M. Watson, Peter C. Hunter, Steve A. McGloughlin, Denis W. Spelman, Adam W. J. Jenney, Kathryn E. Holt

## Abstract

**Background:** *Klebsiella pneumoniae* is a leading cause of extended-spectrum beta-lactamase (ESBL) producing hospital-associated infections, for which elderly patients are at increased risk.

**Methods:** We conducted a 1-year prospective cohort study, in which a third of patients admitted to two geriatric wards in a specialized hospital were recruited and screened for carriage of *K. pneumoniae* by microbiological culture. Clinical isolates were monitored via the hospital laboratory. Colonizing and clinical isolates were subjected to whole genome sequencing and antimicrobial susceptibility testing.

**Results:** *K. pneumoniae* throat carriage prevalence was 4.1%, rectal carriage 10.8% and ESBL carriage 1.7%. *K. pneumoniae* infection incidence was 1.2%. The isolates were diverse, and most patients were colonized or infected with a unique phylogenetic lineage, with no evidence of transmission in the wards. ESBL strains carried *bla*_CTX-M-15 and_ belonged to clones associated with hospital-acquired ESBL infections in other countries (ST29, ST323, ST340).

One also carried the carbapenemase *bla*_IMP-26_. Genomic and epidemiological data provided evidence that ESBL strains were acquired in the referring hospital. Nanopore sequencing also identified strain-to-strain transmission of a *bla*_CTX-M-15_ FIB_K_/FII_K_ plasmid in the referring hospital.

**Conclusions:** The data suggest the major source of *K. pneumoniae* was the patient’s own gut microbiome, but ESBL strains were acquired in the referring hospital. This highlights the importance of the wider hospital network to understanding *K. pneumoniae* risk and infection control. Rectal screening for ESBL organisms upon admission to geriatric wards could help inform patient management and infection control in such facilities.

**Summary:** Patients’ own gut microbiota were the major source of *K. pneumoniae*, but extended-spectrum beta-lactamase strains were acquired in the referring hospital. This highlights the potential for rectal screening, and the importance of the wider hospital network, for local risk management.

## Introduction

*Klebsiella pneumoniae* is an opportunistic bacterial pathogen associated with urinary tract infections (UTIs), pneumonia, septicemia, and wound and soft tissue infections in healthcare settings[1]. One of the ESKAPE pathogens that are collectively responsible for the majority of hard-to-treat infections in hospitalized patients[2] *K. pneumoniae* is frequently multidrug resistant (MDR; defined as resistant to ≥3 classes of antibiotics). Of particular concern are isolates that produce extended-spectrum beta-lactamases (ESBLs) or carbapenemases, which confer resistance to third generation cephalosporins and carbapenems, respectively[2] Among those at risk are infants (who have immature immune systems), and the elderly (who have waning immune defenses), and both are subject to heightened incidence and severity of infections[3]. *K. pneumoniae* carried asymptomatically in the gastrointestinal (GI) tract can disseminate to cause healthcare-associated infections in at-risk individuals[4,5], and we recently reported a positive association between *K. pneumoniae* GI carriage and age amongst patients in an intensive care unit (ICU)[4]. Older age and hospital stays in geriatric and long term care facilities have previously been linked with MDR bacterial colonization and infection[6-8], hence geriatric hospital patients can be considered an at-risk group for carriage and/or infection with *K. pneumoniae*. Here we aimed to investigate the prevalence, diversity and antimicrobial resistance (AMR) of *K. pneumoniae* carried in the GI and respiratory tracts of patients admitted to two geriatric care units.

## Methods

### Ethics

Ethical approval for this study was granted by the Alfred Hospital Ethics Committee (Project number #550/12).

### Recruitment, Specimen and Data Collection

Adult patients aged ≥50 years were recruited from two geriatric medicine wards at the Caulfield Hospital (CH). Verbal consent to participate was required from the patient or an adult responsible for them. Rectal and screening swabs were taken at recruitment, usually within the first week of admission to the ward. Information on age, sex, dates of hospital and ICU admission/s, surgery in the last 30 days, and antibiotic treatment in the last 7 days were extracted from hospital records. All clinical isolates recovered from patients at CH or the referring hospital (Alfred Hospital; AH) patients and identified as *K. pneumoniae* infections by the AH diagnostic laboratory as part of routine care were included in the study. See **Supplementary Methods** for further details.

### Whole genome sequence analysis

DNA was extracted and sequenced via Illumina HiSeq. Multi-locus sequence typing (MLST) was conducted using SRST2 (v2)[9]. Single nucleotide variants (SNVs) were called by aligning reads to a reference genome, and used to infer maximum likelihood phylogenetic trees. Phylogenetic lineages were defined at a threshold of >0.1% divergence. Draft genome assemblies were constructed using SPAdes (v3.6.1) and used to identify capsule loci with Kaptive (v0.4)[10]. Isolates selected for finishing (n=17) were subjected to long read sequencing via Oxford Nanopore MinION and hybrid assembly of long and short reads using Unicycler (v0.4.0)[11] as described previously[12]. All read sets and finished assemblies were deposited in NCBI (accessions in **Table S1)**. See **Supplementary Methods** for full details.

### AMR and plasmid analysis

All clinical and carriage isolates were subjected to antimicrobial susceptibility testing using the Vitek2 GNS card and CLSI breakpoints. AMR genes were identified from Illumina reads using SRST2 (v0.2.0)[9] to screen against the ARG-Annot database[13] (**Table S1**). The location of AMR genes were confirmed by BLAST analysis of finished genome sequences, and the Repository of Antibiotic resistance Cassettes (RAC) database and annotation service[14]. Plasmid incompatibility types and subtypes were identified using PlasmidFinder[15] and established methods for IncC subtyping[16,17].

### Statistical analysis

All statistical analyses were conducted using R (v3.3.1).

## Results

During the one-year study, 296 adults aged ≥50 years and admitted to two geriatric wards of CH (30.5% of 973 patients admitted to these wards) were screened for *K. pneumoniae* carriage via rectal and throat swabs. Participant characteristics are given in **Table 1**. Approximately half (n=144, 49%) were male and age distributions were similar for both genders, with a median age of 84 years (range 55 to 102). Median time of recruitment was day 8 of the current hospital admission (range 1– 83 days). Of the 296 patients screened, 124 (42%) received antimicrobial therapy in the last 7 days, and 34 (11%) underwent one or more surgical procedures in the last 30 days (**Table 1**).

**Table 1.** Characteristics of participants screened for *K. pneumoniae* carriage. Timing is indicated in days of the current admission to hospital (day of admission = day 1). ‘Recent antibiotics’ indicates antimicrobial therapy in the last 7 days prior to screening; ‘Recent surgery’ indicates surgical procedures of any kind in the last 30 days.

|  | **Total** | **Male** | **Female** |
| --- | --- | --- | --- |
| **Number screened** | 296 | 144 (49%) | 152 (51%) |
| **Age** |  |  |  |
| **Median** | 86 | 85 | 86 |
| **Range** | 55 – 102 | 55 – 97 | 58 – 102 |
| **Timing of baseline swab (day of current hospital admission)** |  |  |  |
| **Median** | 8 | 7 | 9 |
| **Range** | 1 – 83 | 1 – 69 | 1 – 83 |
| **Recent antibiotics** | 123 (42%) | 59 (41%) | 64 (42%) |
| **Recent surgery** | 34 (11%) | 15 (10%) | 19 (12%) |
| ***K. pneumoniae* positive culture** |  |  |  |
| **Rectal swab** | 32 (10.8%) | 19 (59%) | 13 (41%) |
| **Throat swab** | 12 (4.1%) | 7 (58%) | 5 (42%) |
| **Both** | 4 (1.4%) | 3 (75%) | 1 (25%) |

### *Klebsiella* carriage

Isolates identified as *K. pneumoniae* were cultured from 13.5% of participants (**Table 1**). We estimate the point prevalence of GI carriage at 10.8% (95% CI, 7.6–15.1%) and throat carriage at 4.1% (95% CI, 2.2–7.2%). The carriage rate was similar among males and females (16% and 11%, respectively; p=0.30 using χ^2^ test). Carriage was not significantly associated with age, sex or day of admission in logistic regression models (**Table S2**), although the study was underpowered to investigate this conclusively. GI carriage of ESBL *K. pneumoniae* was detected in five participants (1.7%), four of these isolates were also MDR (**Table 2**).

**Table 2.**
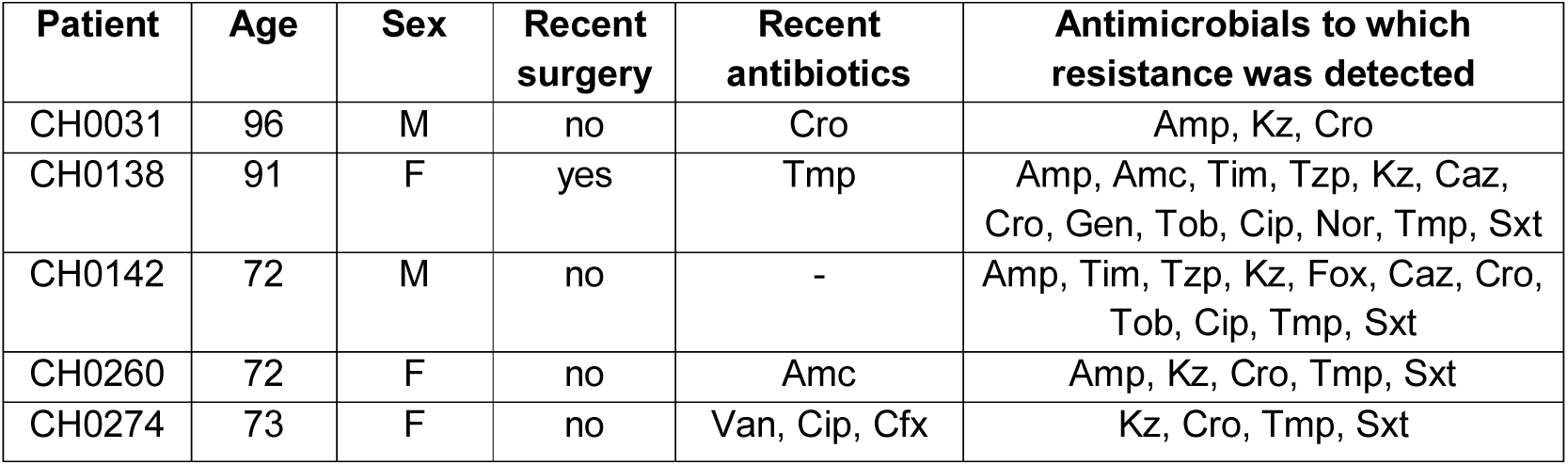
Details of participants with rectal screening swabs positive for ESBL *K. pneumoniae*. CH0031 isolate carried *bla*_CTX-M-55_; the others carried *bla*_CTX-M-15_. ‘Recent antibiotics’ indicates antibiotic treatment in the last 7 days prior to screening; ‘Recent surgery’ indicates surgery of any kind in the last 30 days. AMR phenotyping was conducted using Vitek2 and interpreted using CLSI thresholds, drugs to which resistance was detected are listed as follows: Amp, Ampicillin; Amc, Amoxicillin-clavulanic acid; Tim, Ticarcillin-clavulanic acid; Tzp, Tazobactam-piperacillin; Kz, Cefazolin; Fox, Cefoxitin; Caz, Ceftazidime; Cfx, Cefuroxime; Cro, Ceftriaxone; Gen, Gentamicin; Tob, Tobramycin; Cip, Ciprofloxacin; Nor, Norfloxacin; Tmp, Trimethoprim; Sxt, Trimethoprim-sulfamethoxazole; Van, vancomycin. All four isolates were susceptible to cefepime, meropenem and amikacin.

| Patient | Age | Sex | Recent surgery | Recent antibiotics | Antimicrobials to which resistance was detected |
| --- | --- | --- | --- | --- | --- |
| CH0031 | 96 | M | no | Cro | Amp, Kz, Cro |
| CH0138 | 91 | F | yes | Tmp | Amp, Amc, Tim, Tzp, Kz, Caz, Cro, Gen, Tob, Cip, Nor, Tmp, Sxt |
| CH0142 | 72 | M | no | - | Amp, Tim, Tzp, Kz, Fox, Caz, Cro, Tob, Cip, Tmp, Sxt |
| CH0260 | 72 | F | no | Amc | Amp, Kz, Cro, Tmp, Sxt |
| CH0274 | 73 | F | no | Van, Cip, Cfx | Kz, Cro, Tmp, Sxt |

### *K. pneumoniae* infections

Twelve (1.2%) CH study ward patients (seven female, five male) were diagnosed with *K. pneumoniae* infections (all UTIs). Patient characteristics are shown in **Table 3**. The *K. pneumoniae* UTI rate was very low in non-carriers (0.76%, n=2/264), however the numbers are too small for meaningful comparisons with the rate in carriers (1/32, 3%; note 9 UTIs occurred in patients not screened). Four of the 14 UTI isolates (from 12 patients) were MDR, and three produced ESBLs (**Table 3**).

**Table 3.** Details of *K. pneumoniae* infections (UTI). Characteristics of patients and corresponding urine isolates are provided. Carriage screening swab results are given for patients who were also recruited as participants in the carriage study (NP, not participating). AMR phenotyping was conducted using Vitek2 and interpreted using CLSI thresholds, drugs displaying resistance are listed using the following key: Amp, Ampicillin; Amc, Amoxicillin-clavulanic acid; Tim, Ticarcillin-clavulanic acid; Tzp, Tazobactam-piperacillin; Kz, Cefazolin; Fox, Cefoxitin; Caz, Ceftazidime; Cro, Ceftriaxone; Fep, Cefepime; Mer, Meropenem; Ami, Amikacin; Gen, Gentamicin; Tob, Tobramycin; Cip, Ciprofloxacin; Nor, Norfloxacin; Tmp, Trimethoprim; Sxt, Trimethoprim-sulfamethoxazole. Where two clinical isolates were cultured, these are labelled 1 and 2, and AMR phenotypes are given for both. *=Isolate identified from genome data as *K. variicola*; **=Isolate identified as ESBL producer.

| Patient | Age | Sex | Carriage positive | Antimicrobials to which clinical isolate/s displayed resistance |
| --- | --- | --- | --- | --- |
| CH0110 | 92 | F | no | 1: Amp<br>2: Amp, Tmp and Sxt |
| CH0138 | 91 | F | yes<br>(rectal swab) | 1: Amp, Amc, Tim, Tzp, Kz, Caz, Cro**, Gen, Tob, Cip, Nor, Tmp, Sxt<br>2: Amp, Amc, Tim, Tzp, Kz, Caz, Cro**, Gen, Tob, Cip, Nor, Tmp, Sxt, Fox, Fep, Mer |
| CH0258 | 91 | F | no | Amp, Amc, Tim, Kz, Caz, Cro**, Tob, Tmp, Sxt |
| KC0049 | 78 | F | NP | Amp |
| KC0061 | 90 | F | NP | Amp |
| KC0109 | 69 | M | NP | Amp |
| KC0135 | 88 | F | NP | Amp* |
| KC0136 | 90 | F | NP | Amp* |
| KC0191 | 91 | M | NP | Amp, Amc, Tim, Tzp, Kz, Cro**, Fep, Tob, Tmp, Sxt |
| KC0216 | 86 | M | NP | Amp, Nor |
| KC0302 | 81 | M | NP | Amp |
| KC0303 | 71 | M | NP | Amp |

### Genomic diversity of *Klebsiella*

We sequenced the genomes of all 59 isolates identified as *K. pneumoniae* from patients in the study wards: 12 from throat swabs, 33 rectal swab isolates and 14 UTI isolates, originating from 53 patients. Five carriage isolate genomes were excluded from detailed phylogenetic analysis due to sequence failure (n=1) or mixed culture (n=4). Genome data from the remaining 54 isolates confirmed they were members of the *K. pneumoniae* complex, which are typically indistinguishable biochemically[18]: 37 *K. pneumoniae,* two *Klebsiella quasipneumoniae*, 15 *Klebsiella variicola* (**Figure 1**). UTI isolates were predominantly *K. pneumoniae* (n=12/14, 86%). Carriage isolates were more diverse with only 63% *K. pneumoniae*, however the difference was not statistically significant (p=0.20 using χ^2^ test).

**Figure 1.**
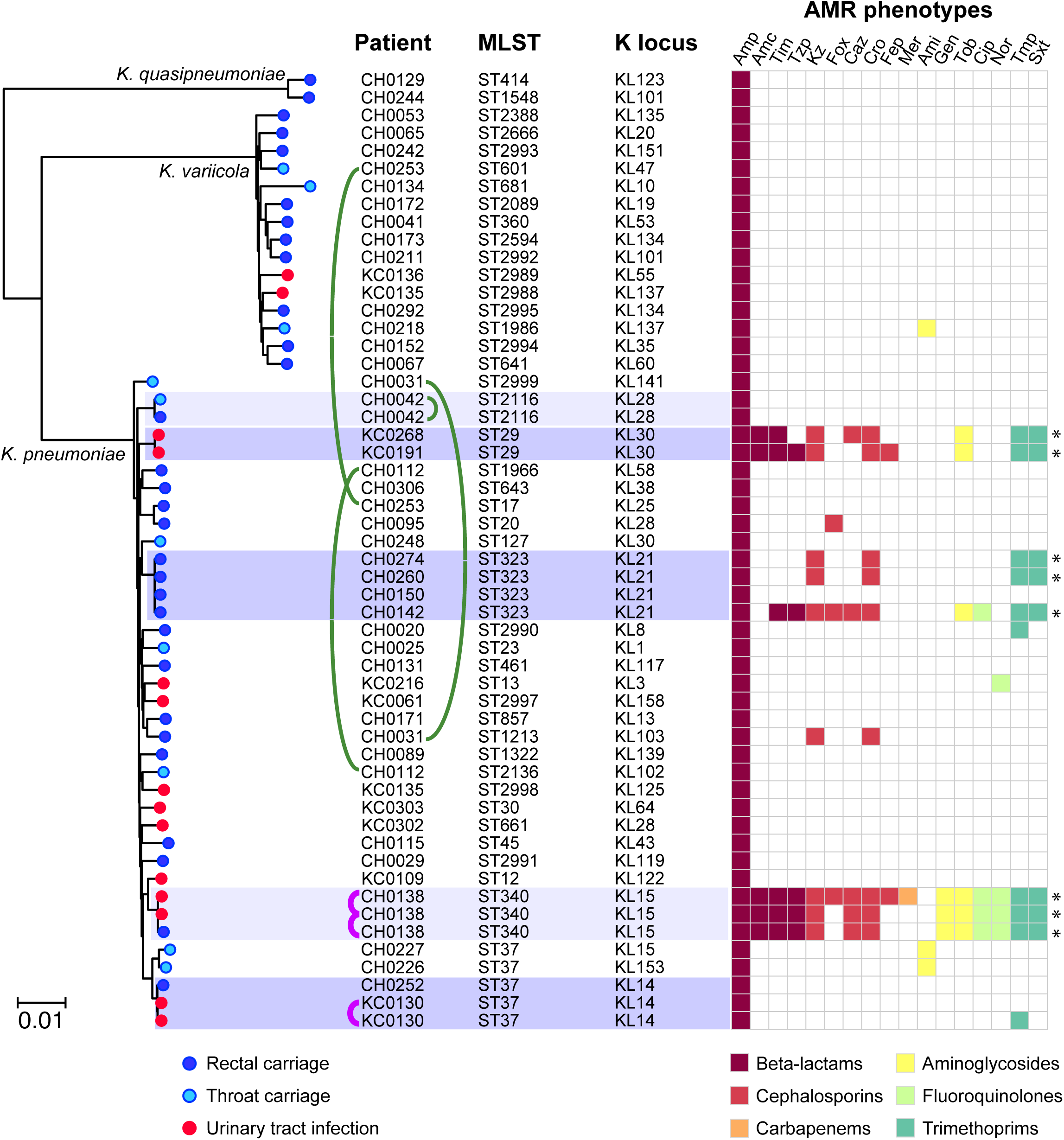
Core genome phylogeny for *Klebsiella* isolated from patients in the study wards, indicating MLST and capsule locus (KL) genotypes, and antimicrobial resistance (AMR) phenotypes. Tree is a maximum likelihood phylogeny inferred from 489,069 SNPs in 3,769 core genes; all branches defining species or lineages (i.e. clades of isolates sharing MLST sequence types) have >90% bootstrap support. Scale bar indicates nucleotide divergence. Clusters of genomes with patristic distance ≤0.1% nucleotide divergence are highlighted. Curved lines indicate isolates from the same patient (green, carriage isolates only; pink, infection isolates). K loci were identified using Kaptive. AMR phenotypes, determined using Vitek2 and interpreted according to CLSI guidelines, are indicated in the heatmap and coloured by drug class according to inset legend; *=multidrug resistance (resistant to ≥3 drug classes). Amp, Ampicillin; Amc, Amoxicillin-clavulanic acid; Tim, Ticarcillin-clavulanic acid; Tzp, Tazobactam-piperacillin; Kz, Cefazolin; Fox, Cefoxitin; Caz, Ceftazidime; Cro, Ceftriaxone; Fep, Cefepime; Mer, Meropenem; Ami, Amikacin; Gen, Gentamicin; Tob, Tobramycin; Cip, Ciprofloxacin; Nor, Norfloxacin; Tmp, Trimethoprim; Sxt, Trimethoprim-sulfamethoxazole.

A core genome tree of the 54 *Klebsiella* isolates (**Figure 1**) showed they represent a diverse population comprising 45 phylogenetically distinct lineages. These included some common MLST sequence types (STs) previously associated with AMR infections in hospitals (ST17, ST20, ST340)[19] or severe community-acquired infections (ST23, ST45)[18,20], as well as 13 novel STs that were submitted to the MLST database for ST assignment (5 *K. pneumoniae* and 8 *K. variicola*). Thirty-seven distinct capsule locus types were also detected[10] (**Figure 1, Table S1**).

Of the four individuals with both rectal and throat isolates, three were colonized with different strains at each site (**Figure 1**). All *K. quasipneumoniae* and *K. variicola* isolates were singleton strains, i.e. each represented a unique lineage detected in a single patient. In contrast, three *K. pneumoniae* lineages were identified in ≥2 patients and thus could potentially indicate transmission in CH (ST323, ST29, ST37; dark shading in **Figure 1**). To investigate further, we used long-read sequencing to generate high quality, completely resolved genome sequences for these isolates (**Methods, Table S3**), and calculated SNV distances between isolates within each cluster (**Table 4**). The ST37 isolates were separated by 1,816 SNVs (433 mutations plus 1,383 SNVs introduced by recombination, **Figure S1**) and thus represent independent strains not linked by recent transmission. The ST323 and ST29 isolates were much more closely related (4–40 SNVs), consistent with recent transmission. These isolates were ESBL and are investigated in detail below.

**Table 4.** Pairwise genetic differences between isolates in multi-patient lineages. Each isolate within a lineage is listed with accompanying patient and sample data, as well as the number of chromosomal and plasmid SNVs detected between the isolate and the first collected isolate from the same lineage. *In INF249, the FIB_K_/FII_K_ plasmid sequence is integrated into the chromosome. NA, no FIB_K_/FII_K_ sequence detected.

| Patient ID | Age | Sex | Sample ID | Specimen | Sample date | SNVs |  |
| --- | --- | --- | --- | --- | --- | --- | --- |
|  |  |  |  |  |  | Chromosome | FIB<sub>K</sub>/FII<sub>K</sub> plasmid |
| ST29 |  |  |  |  |  |  |  |
| KC0191 | 91 | M | INF249 | UTI | 01/12/2013 | - | - |
| CH0258 | 91 | F | INF322 | UTI | 17/02/2014 | 4 | 39* |
| ST323 |  |  |  |  |  |  |  |
| CH0142 | 72 | M | KSB1_4E | Rectal | 10/09/2013 | - | - |
| CH0150 | 85 | M | KSB1_7E | Rectal | 17/09/2013 | 4 | NA |
| CH0260 | 72 | F | KSB1_10J | Rectal | 11/02/2014 | 19 | 0 |
| CH0274 | 73 | F | KSB2_1B | Rectal | 25/02/2014 | 40 | 5 |
| ST37 (KL14) |  |  |  |  |  |  |  |
| CH0110 | 92 | F | INF042 | UTI | 08/05/2013 | - | - |
| CH0110 | 92 | F | INF059 | UTI | 16/05/2013 | 4 | NA |
| CH0252 | 85 | M | KSB1_7J | Rectal | 04/02/2014 | 1,786 | NA |

### MDR mechanisms and transmission

Seven of the 54 isolates were MDR (**Figure 1**). These belonged to the two potential transmission clusters (ST29 and ST323), and UTI and colonizing isolates from patient CH0138 (ST340). To investigate the genetic mechanisms for resistance, we used long-read sequencing to completely resolve these genomes (**Methods**). The ST29 and ST323 MDR isolates each harboured 4–6 plasmids (**Table S3**). Notably, all the acquired AMR genes in these genomes were localized to the same >200 kbp FIB_K_/FII_K_ plasmid backbone encoded conjugative transfer functions and shared close similarity (>90% coverage and 99% identity) with plasmid pKPN3-307_type A from an Italian *K. pneumoniae*[21]. However there were some differences in the MDR regions, resulting in different susceptibility profiles (**Figure S2**).

Both CH ST29 UTI isolates displayed similar AMR profiles and harboured the same set of 12 AMR genes, including the ESBL gene *bla*_CTX-M-15_ (**Figure S2**). In one of the ST29 isolates, INF322, the AMR genes were located on a 243,634 bp circular plasmid (pINF322) carrying the FIB_K_ and FII_K_ replicons.

This entire plasmid sequence was integrated into the chromosome of INF249, within a 23S rRNA gene (**Figure S2A**), flanked by copies of IS*26* and an 8-bp target site duplication (GGCTTTTC). The pINF322 plasmid carried eight copies of IS*26*, and the integration event in INF249 appears to have been mediated by the copy situated next to the *aac6-Ib* gene (**Figure S2A**). The first ST323 carriage isolates, KSB1_4E, carried a plasmid sequence (pKSB1_4E) that differed from pINF322 by just 1 SNV. Two of the other three ST323 isolates carried the identical plasmid backbone to pKSB1_4E (no SNVs). All three harboured *bla*_CTX-M-15_ within a variable MDR region (**Figure S2B**).

We investigated transmission of ST29 and ST323 strains and plasmids based on SNV counts, phylogenetic relationships and hospital admission data (including time in CH and the referring hospital AH). In both the ST29 and ST323 clusters, the data did not support direct transmission between the patients in CH, as in many cases they lacked overlapping time in the wards (**Figure 2A**). Given the close genetic distances between isolates (4–40 SNVs, **Table 4**), we hypothesized that they may be linked by transmission in the referring hospital rather than CH. To investigate this we analysed all ST29 and ST323 clinical isolates identified at AH and CH during the study period, and constructed core genome phylogenies (**Figure 2B**). The CH ST29 isolates belonged to a group of 12 closely related strains (isolated in the last four months of the study) that shared a common ancestor with AH isolates, from which each differed by 1–4 SNVs (**Figure 2B**). Similarly the CH ST323 isolates belonged to a group of 27 strains (isolated throughout the study) that shared a common ancestor with AH isolates from which each differed by between three (early isolates) and 34 SNVs (late isolates).

**Figure 2.**
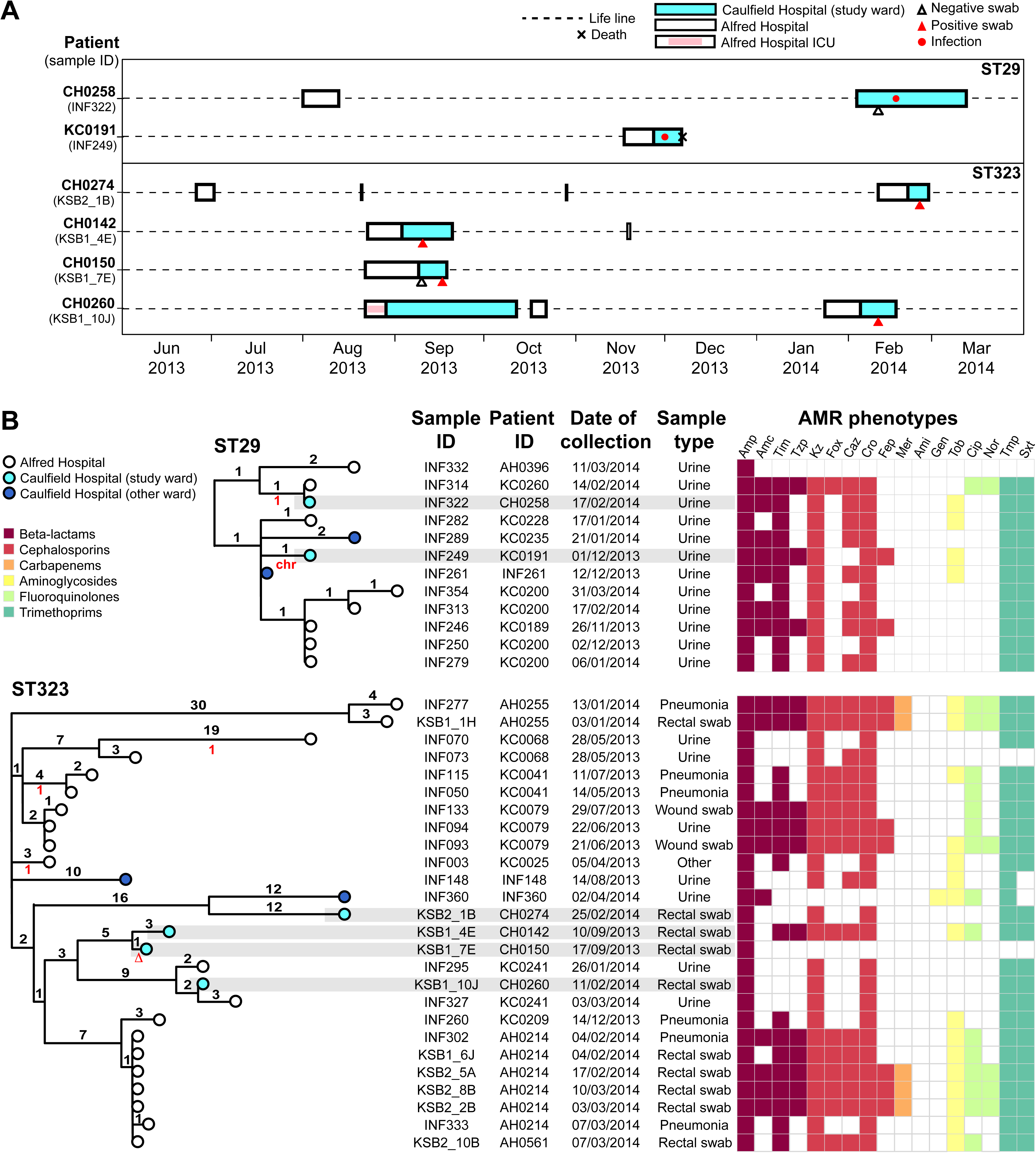
Multidrug resistant ESBL lineages associated with multiple patients in the study wards. **(A)** Timelines for Caulfield and Alfred Hospital stays for patients in the Caulfield Hospital study wards who were infected with ST29 or colonized with ST323 *K. pneumoniae*. (Note ST340 data is shown separately in Figure 3.) Details of all admissions to the Caulfield Hospital (study site) or the Alfred Hospital (referring hospital) during the one-year study period (April 1, 2013 – March 31, 2014) were extracted from the hospital records of these patients. **(B)** Midpoint-rooted core genome phylogenetic trees for all ST29 and ST323 *K. pneumoniae* isolated at Caulfield or Alfred Hospitals during the study period. Tips are coloured by hospital of corresponding specimen collection, according to inset legend. Isolates from the Caulfield Hospital study wards are highlighted in grey. Branches are labelled with the number of chromosomal SNVs defining the branch (black numbers, above branch) and the number of SNVs in the corresponding FIB_K_/FII_K_ plasmid sequence, relative to the major genotype (red numbers, below branch, see Figure S2 for plasmid tree; u=plasmid lost; chr=plasmid integrated into chromosome). Antimicrobial resistance (AMR) phenotypes, determined using Vitek2 and interpreted according to CLSI guidelines, are indicated in the heatmap and coloured by drug class according to inset legend. Amp, Ampicillin; Amc, Amoxicillin-clavulanic acid; Tim, Ticarcillin-clavulanic acid; Tzp, Tazobactam-piperacillin; Kz, Cefazolin; Fox, Cefoxitin; Caz, Ceftazidime; Cro, Ceftriaxone; Fep, Cefepime; Mer, Meropenem; Ami, Amikacin; Gen, Gentamicin; Tob, Tobramycin; Cip, Ciprofloxacin; Nor, Norfloxacin; Tmp, Trimethoprim; Sxt, Trimethoprim-sulfamethoxazole.

All ST29 isolates and n=26/27 ST323 isolates shared the same FIB_K_/FII_K_ plasmid sequence, separated by ≤1 SNV in the backbone sequence. There was variation in the MDR regions resulting in variable AMR phenotypes (**Figures 2C, S2C**), but *bla*_CTX-M-15_ was retained in all but two isolates.

### Carbapenem resistance

Meropenem resistant *K. pneumoniae* was detected in one CH patient (CH0138). Their rectal carriage isolate and two clinical urine isolates, collected on the same day, shared near-identical chromosomal sequences separated by a single SNV (**Figure 3**). Patient CH0138 was discharged after two months and re-admitted to AH nearly two months later. *K. pneumoniae* was isolated from clinical urine samples on days four and ten of the second admission. These were also ST340, separated from the earlier isolates by 3–5 SNVs (**Figure 3**).

**Figure 3.**
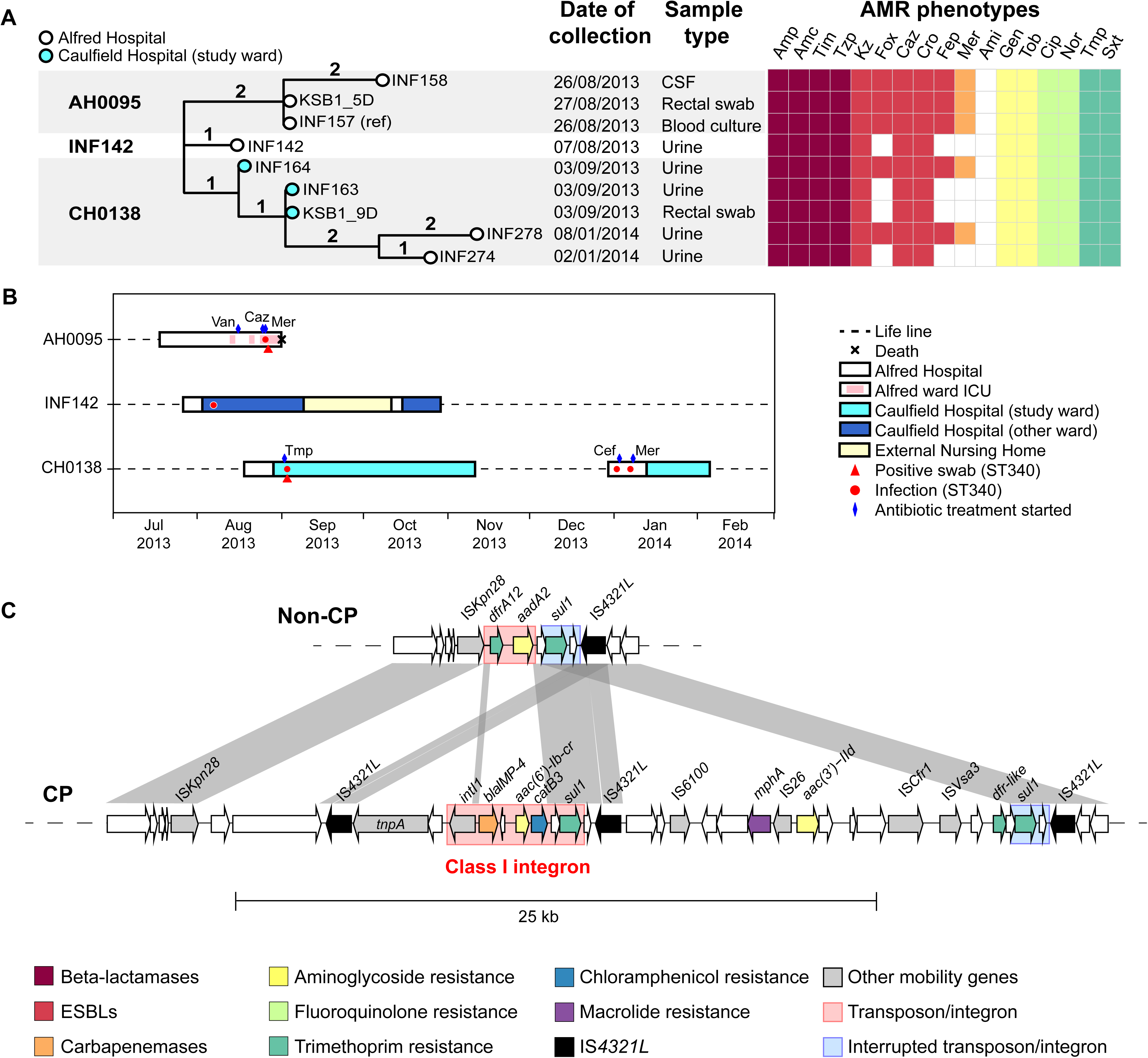
Carbapenemase lineage ST340. **(A)** Core genome phylogenetic tree of all ST340 isolates identified at Caulfield or Alfred Hospitals during the study period. Branches are labelled with number of chromosomal SNVs. CSF, cerebral spinal fluid. Antimicrobial resistance (AMR) phenotypes, determined using Vitek2 and interpreted according to CLSI guidelines, are indicated in the heatmap and coloured by drug class according to inset legend. Amp, Ampicillin; Amc, Amoxicillin-clavulanic acid; Tim, Ticarcillin-clavulanic acid; Tzp, Tazobactam-piperacillin; Kz, Cefazolin; Fox, Cefoxitin; Caz, Ceftazidime; Cro, Ceftriaxone; Fep, Cefepime; Mer, Meropenem; Ami, Amikacin; Gen, Gentamicin; Tob, Tobramycin; Cip, Ciprofloxacin; Nor, Norfloxacin; Tmp, Trimethoprim; Sxt, Trimethoprim-sulfamethoxazole. **(B)** Timelines for hospital stays; all admissions to the Caulfield Hospital (where the study wards were located) or the Alfred Hospital (referring hospital) during the 1-year study period (April 1, 2013 – March 31, 2014) were extracted from hospital records of these patients. **(C)** Comparison of AMR regions in the two forms of IncC plasmid identified in these isolates; AMR and transposase genes are labelled and coloured according to the inset legend. CP=carbapenemase producing, defined by presence of *bla*IMP-4 gene and phenotypic meropenem resistance (Mer in panel **A**).

Two distinct AMR patterns were observed amongst the CH0138 isolates: all were ESBL and MDR, however the first and last urine isolates were additionally resistant to cefoxitin, cefepime and meropenem (**Figure 3A**).

Completion of the genomes with long reads showed the MDR phenotypes were conferred by (a) chromosomal insertions of *sul1*, *aadA2* (via a type 1 integron) and *bla*_CTX-M-15_ (via IS*Ecp1*); (b) two large conjugative plasmids: a novel 90 kbp plasmid sharing two replicon genes (*repB* and *repE*) with pK245 (accession DQ449578), carrying *aac3-IIa, aac6Ib-cr, catB4, strAB, bla_OXA-1_, sul2* and *qnrB1*; and (c) an IncC type 2 plasmid[16] (also known as IncA/C2 plasmid sequence type 3[17]) carrying *bla_TEM_, aadA2, tetA, dfrA14* (**Figure 3C, Table S4**). In the meropenem-resistant strains, the IncC plasmid also carried the carbapenemase *bla*_IMP-26_ as well as *mphA, aac3-IIa* and *catB3* (**Figure 3C**).

Comparison to clinical isolates identified in AH during the study period identified four additional ST340 isolates from two patients, which shared the same plasmids and chromosomal AMR determinants and were separated from the first CH0138 isolate by 2–5 chromosomal SNVs (**Figure 3A, Table S4**). Three strains (blood, CSF and rectal swab) were isolated from patient AH0095 on admission to ICU with sepsis. This followed a month-long stay in AH, which overlapped with the AH admissions of the other two patients, likely providing the opportunity for transmission (**Figure 3B**).

## Discussion

We estimated overall carriage prevalence in the first week of admission to the geriatric unit at 13.5%, with 10.8% GI carriage and 4% throat carriage. GI carriage prevalence was significantly higher than we estimated previously for patients admitted to the AH ICU with no recent healthcare exposure[4] (5.9%, OR 1.94 [95% CI, 1.0–3.7], p=0.03) but significantly lower than that for ICU patients who had been in hospital for >2 days or had recent surgery (19%, OR 0.52 [0.3–0.9], p=0.02). The rate was also lower than that recently estimated for ICU and hematology/oncology patients at a US hospital[5] (23%, OR 0.41 [0.27–0.60], p=7x10^-7^).

Colonizing isolates were diverse (**Figure 1**) and mostly unique to individual participants, consistent with our earlier observations in ICU patients[4] and the recent US study[5]. Hence in most cases, these strains likely represent established members of the patients’ microbiota rather than hospital-acquired bacteria. In the ICU, *K. pneumoniae* carriage is significantly associated with subsequent infection (OR ≥4)[4,5]. In the current setting, *K. pneumoniae* infections occurred at a similar rate (1.2% vs 1.8–2.2%, p>0.1) but the number of cases was too small to explore the link between colonization and infection directly. Notably the 12 patients with *K. pneumoniae* infections were infected with 11 different lineages, consistent with most patients developing a UTI from their own microbiota. The exceptions were two UTIs associated with ESBL ST29, acquired by both patients in the referring hospital (**Figure 2**).

ESBL carriage (1.7%) was much rarer than previously reported in point prevalence studies of geriatric units or long term care facilities[22,23]. This is unsurprising as we aimed to screen patients during the first week of their CH stay and only focused on *K. pneumoniae*. The MDR strains identified at CH all belonged to lineages that have been previously associated with hospital outbreaks of ESBL and/or carbapenemase producing (CP) *K. pneumoniae* on other continents (ST29, ST323, ST340)[24],[25], indicating the emergence of globally distributed ESBL strains in Australia. The spread of internationally common ST258 strains harbouring the *K. pneumoniae* carbapenemase (KPC), which are common internationally, have also been recently detected in Australia[26]. FIB_K_/FII_K_ plasmids are frequently reported as disseminators of *bla*_CTX-M-15_ and other AMR genes in *K. pneumoniae*, *E. coli* and other enteric bacteria isolated from humans, animals and the environment[27,28]. The IncC plasmid carrying *bla*_IMP_ was recently reported in *K. pneumoniae* (ST unknown) isolated from Australian wild birds[29], indicating that this plasmid and possibly the host strain are involved in spreading carbapenemases between animals and humans.

This study employed Illumina short-read WGS to identify high-confidence SNVs with which to identify lineages and AMR genes, bolstered by long-read sequencing to resolve plasmids and AMR gene context and to maximize resolution for detecting transmission. While this strategy has been used in other studies[30-33], ours is the first to adopt multiplex nanopore sequencing and hybrid assembly to rapidly and cost-efficiently complete genome sequences of interest in a high-throughput manner[11,12] (**Tables S3, S4**).

However, unravelling the source of MDR *K. pneumoniae* in CH also required exploring patient movement prior to CH admission, and phylogenetic context provided by additional isolates from the referring hospital.

Our analyses revealed that all MDR *K. pneumoniae* isolated at CH were linked to transmission clusters at the referring hospital (**Figures 2-3**). This suggests that MDR carriage is rare in the community, but MDR ESBL *K. pneumoniae* were occasionally acquired in the referring hospital prior to transfer to CH (ESBL GI carriage prevalence 1.7%, ESBL infection incidence 0.31%). These findings are likely generalisable to other hospital referral networks and highlight the benefits of exploring transmission at a multi-facility level[34,35]. However, larger studies will be needed to confirm the importance of referral networks for transmission of ESBL organisms, as has been demonstrated for MRSA and *Clostridium difficile*[34,35], and to explore specific risk factors and the relevance to healthcare-associated infections.

It is noteworthy that the two ESBL lineages transmitting at AH (ST323 and ST29) shared the same FII_K_/FIB_K_ *bla*_CTX-M-15_ plasmid. Given that ST323 appears to have been circulating at AH for months prior to the common ancestor of the ST29 strains, we hypothesise that the plasmid transferred from ST323 to ST29 within the hospital, promoting transmission of ST29 within AH. This highlights the importance of tracking ESBL or CP plasmids as well as their host strains, as has been noted previously[30,31,36]

The spread of CP *K. pneumoniae* within the referring hospital is concerning. The CP ST340 strain carrying *bla*_IMP_ and *bla*_CTX-M-15_ was identified in clinical isolates from two patients, and additional variants lacking *bla*_IMP_ were found in one of these patients plus a ST340 UTI isolate from a third patient (**Figure 3**). Notably the ST340 strain was detected on rectal swabs from both patients who were screened for carriage (one CH patient in the present study; one ICU patient in the prior study[4]). However, it is encouraging to note that we detected no evidence of its transmission within the CH wards.

Only one third of patients were recruited for GI carriage screening at CH. While two of the three patients with MDR infections were not screened, given the strong evidence that they acquired their infecting strain (ESBL ST29) in the referring hospital (**Figure 2**) it is likely that they would have been detected as ESBL carriers if they had been swabbed on arrival. Surveillance swabs are frequently recommended to screen for carriage of MDR organisms in a variety of settings[37-39], but their role in the management of ESBL or CP Gram negative infections outside of outbreaks remains controversial[40]. Our study suggests that screening upon transfer from tertiary referral hospitals to specialized hospitals could be valuable for management or prevention of MDR infections.

## Acknowledgments

The authors gratefully acknowledge the contribution and support of Janine Roney, Mellissa Bryant, Jennifer Williams, Iain Abbott and Noelene Browne at the Alfred Hospital, and the sequencing team at the Wellcome Trust Sanger Institute. We also thank the team of curators at the Institut Pasteur MLST system (Paris, France) for importing novel alleles, profiles and/or isolates at http://bigsdb.web.pasteur.fr. The authors declare no conflicts of interest.

## References

1. Podschun R, Ullmann U. Klebsiella spp. as nosocomial pathogens: epidemiology, taxonomy, typing methods, and pathogenicity factors. Clin Microbiol Rev 1998; 11:589–603.

2. Pendleton JN, Gorman SP, Gilmore BF. Clinical relevance of the ESKAPE pathogens. Expert Rev Anti Infect Ther 2013; 11:297–308.

3. McElhaney JE, Effros RB. Immunosenescence: what does it mean to health outcomes in older adults? Curr Opin Immunol 2009; 21:418–424.

4. Gorrie CL, Mirèeta M, Wick RR, et al. Gastrointestinal carriage is a major reservoir of Klebsiella pneumoniae infection in intensive care patients. Clin Infect Dis 2017; 65:208–215.

5. Martin RM, Cao J, Brisse S, et al. Molecular Epidemiology of Colonizing and Infecting Isolates of Klebsiella pneumoniae. mSphere 2016; 1:e00261–16.

6. Venkatachalam I, Yang HL, Fisher D, et al. Multidrug-Resistant Gram-Negative Bloodstream Infections among Residents of Long-Term Care Facilities. Infect Control Hosp Epidemiol 2014; 35:519–526.

7. Denkinger CM, Grant AD, Denkinger M, Gautam S, DAgata EMC. Increased multi-drug resistance among the elderly on admission to the hospital–A 12-year surveillance study. Arch Gerontol Geriatr 2013; 56:227–230.

8. Gruber I, Heudorf U, Werner G, et al. Multidrug-resistant bacteria in geriatric clinics, nursing homes, and ambulant care – Prevalence and risk factors. Int J Med Microbiol 2013; 303:405–409.

9. Inouye M, Dashnow H, Raven L-A, et al. SRST2: Rapid genomic surveillance for public health and hospital microbiology labs. Genome Med 2014; 6:90.

10. Wyres KL, Wick RR, Gorrie CL, et al. Identification of Klebsiella capsule synthesis loci from whole genome data. Microb Genom 2016; 2. doi:10.1099/mgen.0.000102

11. Wick RR, Judd LM, Gorrie CL, Holt KE. Unicycler: Resolving bacterial genome assemblies from short and long sequencing reads. PLoS Comput Biol 2017; 13:e1005595.

12. Wick RR, Judd LM, Gorrie CL, Holt KE. Completing bacterial genome assemblies with multiplex MinION sequencing. Microb Genom 2017; 3. doi:10.1099/mgen.0.000132

13. Gupta SK, Padmanabhan BR, Diene SM, et al. ARG-ANNOT, a new bioinformatic tool to discover antibiotic resistance genes in bacterial genomes. Antimicrob Agents Chemother 2014; 58:212–220.

14. Tsafnat G, Copty J, Partridge SR. RAC: repository of antibiotic resistance cassettes. Database 2011; 2011.

15. Carattoli A, Zankari E, García-Fernández A, et al. In silico detection and typing of plasmids using PlasmidFinder and plasmid multilocus sequence typing. Antimicrob Agents Chemother 2014; 58:3895–3903.

16. Harmer CJ, Hall RM. The A to Z of A/C plasmids. Plasmid 2015; 80:63–82.

17. Hancock SJ, Phan M-D, Peters KM, et al. Identification of IncA/C Plasmid Replication and Maintenance Genes and Development of a Plasmid Multilocus Sequence Typing Scheme. Antimicrob Agents Chemother 2017; 61.

18. Holt KE, Wertheim H, Zadoks RN, et al. Genomic analysis of diversity, population structure, virulence, and antimicrobial resistance in Klebsiella pneumoniae, an urgent threat to public health. Proc Natl Acad Sci U S A 2015; 112:E3574–E3581.

19. Wyres KL, Holt KE. Klebsiella pneumoniae population genomics and antimicrobial-resistant clones. Trends Microbiol 2016; 24:944–956.

20. Bialek-Davenet S, Criscuolo A, Ailloud F, et al. Genomic Definition of Hypervirulent and Multidrug-Resistant Klebsiella pneumoniae Clonal Groups. Emerg Infect Dis 2014; 20:1812–1820.

21. Villa L, Feudi C, Fortini D, et al. Diversity, virulence, and antimicrobial resistance of the KPC-producing Klebsiella pneumoniae ST307 clone. Microb Genom 2017; 3. doi:10.1099/mgen.0.000110.

22. March A, Aschbacher R, Dhanji H, et al. Colonization of residents and staff of a long-term-care facility and adjacent acute-care hospital geriatric unit by multiresistant bacteria. Clin Microbiol Infect 2010; 16:934–944.

23. Lim CJ, Cheng AC, Kennon J, et al. Prevalence of multidrug-resistant organisms and risk factors for carriage in long-term care facilities: a nested case-control study. J Antimicrob Chemother 2014; 69:1972–1980.

24. Mansour W, Grami R, Ben Haj Khalifa A, et al. Dissemination of multidrug-resistant blaCTX-M-15/IncFIIk plasmids in Klebsiella pneumoniae isolates from hospital- and community-acquired human infections in Tunisia. Diagn Microbiol Infect Dis 2015; 83:298–304.

25. Dolejska M, Brhelova E, Dobiasova H, et al. Dissemination of IncFIIKtype plasmids in multiresistant CTX-M-15-producing Enterobacteriaceae isolates from children in hospital paediatric oncology wards. Int J Antimicrob Agents 2012; 40:510–515.

26. Kwong JC, Lane C, Romanes F, et al. Real-time genomic and epidemiological investigation of a multi-institution outbreak of KPC-producing Enterobacteriaceae: a translational study. bioRxiv 2017. doi:10.1101/175950

27. Mathers AJ, Peirano G, Pitout JDD. The role of epidemic resistance plasmids and international high-risk clones in the spread of multidrugresistant Enterobacteriaceae. Clin Microbiol Rev 2015; 28:565–591.

28. Lahlaoui H, Ben Haj Khalifa A, Ben Moussa M. Epidemiology of Enterobacteriaceae producing CTX-M type extended spectrum β-lactamase (ESBL). Med Mal Infect 2014; 44:400–404.

29. Papagiannitsis CC, Kutilova I, Medvecky M, Hrabak J, Dolejska M. Characterization of the Complete Nucleotide Sequences of IncA/C2 Plasmids Carrying In809-Like Integrons from Enterobacteriaceae Isolates of Wildlife Origin. Antimicrob Agents Chemother 2017; 61.

30. Conlan S, Park M, Deming C, et al. Plasmid Dynamics in KPC-Positive Klebsiella pneumoniae during Long-Term Patient Colonization. mBio 2016; 7.

31. Mathers AJ, Cox HL, Kitchel B, et al. Molecular dissection of an outbreak of carbapenem-resistant enterobacteriaceae reveals Intergenus KPC carbapenemase transmission through a promiscuous plasmid. mBio 2011; 2:e00204–e00211.

32. Stoesser N, Giess A, Batty EM, et al. Genome sequencing of an extended series of NDM-producing Klebsiella pneumoniae isolates from neonatal infections in a Nepali hospital characterizes the extent of community-versus hospital-associated transmission in an endemic setting. Antimicrob Agents Chemother 2014; 58:7347–7357.

33. Martin J, Phan HTT, Findlay J, et al. Covert dissemination of carbapenemase-producing Klebsiella pneumoniae (KPC) in a successfully controlled outbreak: long- and short-read whole-genome sequencing demonstrate multiple genetic modes of transmission. J Antimicrob Chemother 2017; :dkx264–dkx264.

34. Donker T, Wallinga J, Slack R, Grundmann H. Hospital Networks and the Dispersal of Hospital-Acquired Pathogens by Patient Transfer. PLoS ONE 2012; 7:e35002.

35. Simmering JE, Polgreen LA, Campbell DR, Cavanaugh JE, Polgreen PM. Hospital Transfer Network Structure as a Risk Factor for Clostridium difficile Infection. Infect Control Hosp Epidemiol 2015; 36:1031–1037.

36. Sheppard AE, Stoesser N, Wilson DJ, et al. Nested Russian Doll-Like Genetic Mobility Drives Rapid Dissemination of the Carbapenem Resistance Gene blaKPC. Antimicrob Agents Chemother 2016; 60:3767–3778.

37. Fernández J, Bert F, Nicolas-Chanoine M-H. The challenges of multidrug-resistance in hepatology. J Hepatol 2016; 65:1043–1054.

38. Humphreys H. Controlling the spread of vancomycin-resistant enterococci. Is active screening worthwhile? J Hosp Infect 2014; 88:191–198.

39. Humphreys H, Becker K, Dohmen PM, et al. Staphylococcus aureus and surgical site infections: benefits of screening and decolonization before surgery. J Hosp Infect 2016; 94:295–304.

40. Otter JA, Mutters NT, Tacconelli E, Gikas A, Holmes AH. Controversies in guidelines for the control of multidrug-resistant Gram-negative bacteria in EU countries. Clin Microbiol Infect 2015; 21:1057–1066.

